# Controlling Molecular Transport through Nanopores by Dynamic Aperture Sizing

**DOI:** 10.64898/2026.09.01.748496

**Authors:** Justin N. Cronk, Alessio Angeli Bufalini, Morteza Aramesh

## Abstract

Molecular transport through a nanopore determines the information that can be recovered from a translocation signal, yet it remains difficult to control in conventional solid-state nanopores. Rapid translocation reduces the information content and fixed nanopore geometries limit the dimensionality of the signal. Here, we control molecular transport through the development of the pipette–elastomer interfacial nanopore (PEIN), a dynamically reconfigurable solid-state nanopore which addresses these limitations. A PEIN is formed by depressing a glass nanopipette into a soft elastomer, progressively constricting its aperture and enabling continuous control over aperture size, while retaining the simplicity and favourable noise characteristics of glass nanopipette sensing. Using the dynamic aperture size control, DNA velocities could be controlled over more than a twofold range, with dwell times two orders of magnitude greater than observed in glass nanopipettes. DNA-origami rulers further revealed a progressive reduction in polymer velocity during translocation, indicating that hydrodynamic drag alone is insufficient to model forces on the DNA polymer. Finally, by using single- and double-stranded DNA and gold nanoparticles as molecular standards, we demonstrate reversible, size-selective molecular gating with sub-nanometre control. These results establish the PEIN as an accessible platform for controlling molecular transport and probing the relationships among biopolymer conformation, nanoscale confinement and translocation dynamics.

## Introduction

The simplicity of nanopore sensing, that the structural information of a passing molecule can be deduced from the ionic current modulation it causes, has allowed it to become a powerful and commercialised single-molecule microscopy technique.^1,2^ However, the sequence, structure and conformation of a molecule can only be inferred from a nanopore signal if its transport through the aperture occurs on an informative timescale and, for conformation and structure, without denaturing the biopolymer. Therefore, control of molecular transport is necessary as it is not merely a consequence of nanopore sensing, but one of its principal determinants. Biological nanopores illustrate this since, while single-stranded (ss)DNA can freely translocate through biological nanopores,^3,4^ sequence-resolved readout relies on enzyme ratcheting.^5–7^ Similarly, the lack of residue-resolution enzymatic motors for transport of amino-acid chains is one of the key limitations preventing their *de novo* sequencing,^8^ with current approaches such as ClpX falling short.^9^ A further limitation of the biological nanopores used for sequencing is their apertures, which are typically too small for folded biopolymers to translocate (1.4 nm in *α*-haemolysin),^10^ and therefore transport sacrifices information on a biopolymer’s structure. Yet these structural features are central to the function of proteins and RNAs, which depend on both precise folding and dynamic conformational rearrangements.

Conversely, solid-state nanopores can be fabricated with much broader control over pore geometry and can thus interrogate folded, partially folded and globular proteins under non-denaturing conditions, as well as higher-order conformations, unfolding and stretching in the pore’s electric field and non-conventional DNA forms.^11–15^ Nevertheless, molecular transport is more difficult to control in solid-state nanopores and thus, while more aperture-size diverse, short dwell times and fabrication variability have so far precluded routine sequence-resolved readout.^16,17^ In spite of these challenges, measurements of properties including the shape, volume, charge, rotational diffusion coefficient and dipole moment of biopolymers are possible.^18^ Strategies to slow transport have therefore included reducing aperture size to increase analyte-pore interactions and engineering the pore surface chemistry.^13,19,20^

Interfacial nanopores provide an alternative means of controlling transport by allowing the aperture itself to be adjusted *in situ* through the occlusion of a solid-state nanopore upon contact with and depression of an elastic surface. Introducing dynamic aperture size as a new empirical dimension has allowed the study of fibronectin secretion,^21^ localised recordings next to single neurons,^22^ and generally detection of various biomolecules on functionalised surfaces.^23^ Although these results demonstrate the versatility of dynamically formed apertures, they were achieved using piezo-controlled atomic force microscope (AFM) interfacial nanopores which require specialised instrumentation with limited throughput. Moreover, the system did not demonstrate unequivocal control of transport over either molecular dwell time or conformation. Nonetheless, the promise of dynamic control of molecular transport through solid-state nanopores warrants the development and more rigorous characterisation of an alternative interfacial nanopore architecture.

Given the remarkable ability of glass nanopipettes to sense conformational changes in DNA and RNA structures, despite high translocation speeds,^24,25^ they offered a natural platform to develop such an interfacial nanopore. In this article, we present the pipette-elastomer interfacial nanopore (PEIN) in order to control molecular transport. Formed by the constriction of a glass nanopipette by a soft elastomer as it is depressed by a micromanipulator, this solid-state nanopore has the *in-situ* aperture size control of an interfacial nanopore, while maintaining much of the simplicity of glass nanopipette sensing. We characterise the electrical and mechanical properties of a PEIN and its formation, showing that the increased complexity of the pore does not detrimentally affect noise characteristics or translocation dynamics. We further characterise a polymer’s velocity profile in the PEIN using double-stranded (ds)DNA and DNA origami rulers, showing a significant increase in molecular dwell times and quantifying the relationship between DNA dwell time and inferred aperture size. Finally, we demonstrate size-selective molecular gating by dynamically limiting the aperture to permit the free translocation of only molecules smaller than the aperture. Together, these measurements demonstrate the potential of the PEIN as a dynamically reconfigurable solid-state nanopore for controlling molecular transport and probing the relationship between biopolymer conformation, confinement and translocation dynamics.

## Results

### Mechanical Gating of Ionic Transport

The working principle of a PEIN is analogous to other nanopores: the current between two solutions (typically high salt) separated by a nanopore-containing membrane is measured when a voltage is applied across. We form a PEIN *in situ* at the interface of a polydimethylsiloxane (PDMS) coated Petri dish and the aperture of a filament-pulled glass nanopipette of ~ 300 nm (Figure 1a). This is achieved by mounting the pipette on a micromanipulator at a working angle of *θ* (between 30 and 45°) and lowering into the PDMS surface. As the PDMS deforms around the pipette, the effective aperture; the narrowest constriction, shrinks. This process is dynamic and temporally limited by the relaxation time of the PDMS and response time of the micromanipulator (in this instance ~ 250 ms) and, surprisingly for the 300 nm glass pipettes used, does not result in failure or fracture of the nanopipette (verified by repeated formation and withdrawal and monitoring of the current response, Figure SI-D1). We developed custom software which employs a proportional-integral controller to maintain a target resistance through a feedback loop whereby the height is adjusted in response to current changes: withdrawal from the surface opens up the aperture, increasing the measured current and *vice versa* (contrary to previous dynamic nanopores, which were force-controlled AFMs).^21–23^ This software also handles data acquisition, event detection and voltage control. While aperture size is tuned by current feedback, optical access to the dish is maintained, and therefore the deformation can be directly observed. The entire nanopore system is vibrationally isolated above a weighted optical table.

**Figure 1:**
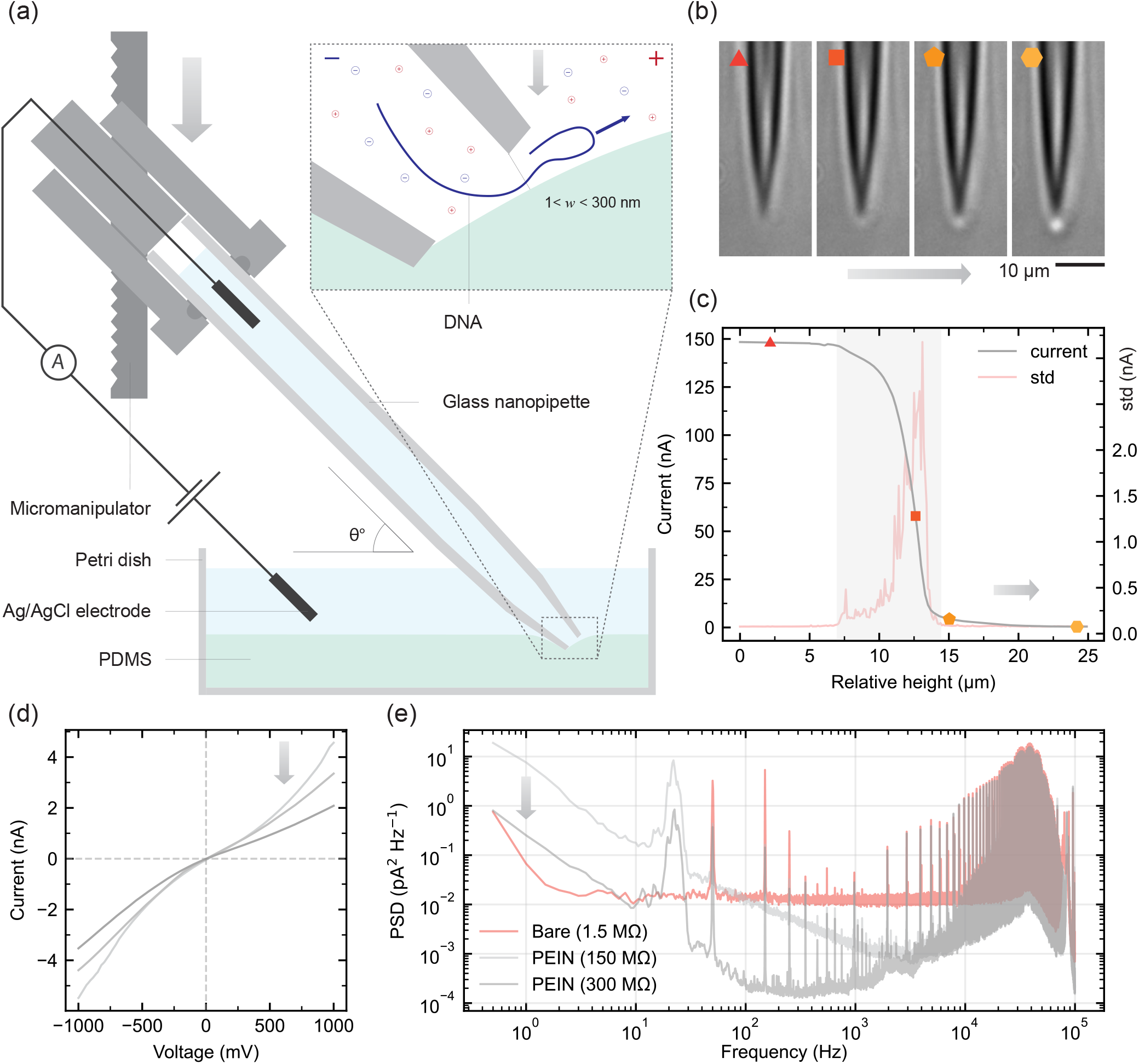
(a) Schematic of a pipette-elastomer interfacial nanopore (PEIN), showing a glass nanopipette depressing a polydimethylsiloxane (PDMS) layer by micromanipulation in the negative *z* direction, with a DNA polymer translocating the effective nanopore (narrowest constriction). Arrows illustrate depression, correlated with effect in (b)-(e).(b) Differential Interference Contrast (DIC) images of nanopipette lowered onto PDMS surface, with increased polygon order indicating further depression, visible as increased brightness at the tip (implies greater curvature and thus bending of the PDMS). The second image corresponds to the first observation of PDMS deformation. (c) Current response to 200 mV applied across the nanopore in 2 M KCl as it approaches the surface with the standard deviation (std) in current shown in red. The zero point in height is defined as 99% of *I*_0_, with height measured from the micromanipulator. The overlaid polygons correspond to the DIC images in (b). The shaded region is the pore-forming region of the approach with high std. (d) Current-voltage (I-V) curves for a PEIN in 2 M KCl solution, with darker shading corresponding to depression in the *z* direction, showing increased resistance (smaller aperture size). (e) Power spectral density (PSD) from 100 kHz bandwidth recordings of a bare nanopipette and PEINs at two resistances.

In order to properly contextualise the PEIN, an understanding of the manner in which the initial bare nanopipette’s aperture is occluded is paramount. As mentioned, optical access is maintained, allowing the pipette approach and subsequent deformation of the PDMS to be imaged. This was achieved using differential interference contrast (DIC) microscopy, which then allows for curvature of the otherwise transparent PDMS to be imaged (Figure 1b). These microscope images show how the PDMS deforms around the approaching pipette as it is pressed further into its surface, increasing the measured resistance and correspondingly reducing the aperture size. These images can then be correlated to approach curves (Figure 1c; 2 M KCl, 200 mV): the current response to the height change as the pipette is lowered into the surface. The zero point is defined as the point at which the current has fallen to 99% of its original value, *I*_0_.

Apart from the oblique pipette angle, the approach of a PEIN pipette is analogous to that of a scanning ion conductance microscopy (SICM) pipette. Therefore, we can make quantitative conclusions about the contact point in our curve. However, SICM usage requires a maximum of only a few percentage points in current reduction, and therefore we must extrapolate from literature to the contact point. Far from the surface, the access resistance at an acute angle is equivalent to a normal pipette and accordingly the distance travelled from 99% to 98% of *I*_0_ (roughly 2.5 *µ*m) is the same order of magnitude as literature for a high conical angle, large aperture pipette.^26^ Corroborating this, no deformation of the PDMS is observed in the DIC imaging, and indeed, with the surface and focal plane being coincident, the pipette remains out of focus, indicating it remains above (Figure 1b; triangle). A normal pipette would then approach zero current within 1-10 *µ*m as the access resistance diverges (ignoring current leakage).^27^ Naturally this is extended across several micrometres in PEINs as even up to the contact point, the aperture remains largely open. Furthermore, the height is measured at the micromanipulator, but bending of the glass capillary following contact with the surface is such that the actual *z*-depression at the tip will be smaller. From microscopy, we assign the contact point to the first sign of PDMS deformation, which is then marked on the approach curve (red square, Figure 1c). This aligns with the standard deviation peak and inflection point of the curve, with the latter indicating the surface is deformed around the tip on further height reductions, yielding relatively reduced changes in current. The broad and high magnitude peak in the standard deviation plot is indicative of the proportionally large change in access resistance with height close to the contact point and thus out-of-phase vibrations here can then cause large changes in resistance and correspondingly aperture size.

As the pipette is lowered further, the PDMS begins to deform around it, with the resulting geometry influenced by the initial size of the aperture. We model the deformation using finite element methods (FEM) (Figure SI-A1), indicating the effective nanopore aperture to be elliptical. From this we can infer that should too large pipettes be used, the PEIN becomes too slit-like in geometry, harming signal-to-noise characteristics. Conversely, too small a pipette risks puncturing the PDMS or breaking the tip when depressed. PDMS puncturing also caused the only instances observed of nanopore clogging; although, given that only DNA polymers were used in this work, this is unsurprising. Illustrative DIC images of puncturing are included in Figure SI-A2 for reference.

As the pipette depresses into the PDMS, the two become mechanically coupled, reducing the influence of vibrations. It is in this ‘stable’ regime (pentagon to hexagon in Figure 1b & c) that single-molecule sensitivity is achieved and our measurements are conducted. In this region, the height can be adjusted over more than 10 *µ*m for resistances between ~ 50 MΩ and ~ 400 MΩ, providing high aperture size resolution, and is sufficient to cover a broad range of molecular sizes, which we show later. This resolution is finer than the 20 nm step-size micromanipulator used in this work due to both the deformation of the PDMS and to bending by the glass capillary.

The electrochemical response of the nanopore was characterised in the stable regime by generating current-voltage (I-V) plots in 2 M KCl 1*×* TE (10 mM Tris-HCl, 1 mM EDTA, pH 9.0) (Figure 1d). These are non-linear and show some ionic current rectification (ICR) not seen in the I-V curve of the bare nanopipette. Rectification can be quantified as the ratio of currents at opposite polarities: *r*(*V*) = |*I*(*V*_+_)*/I*(*V*_*−*_) |; therefore, at 1000 mV in 2 M KCl, a PEIN has a ratio, *r*(1), of ~ 1.5. Although unexpected at such high salt concentrations, ICR arising from asymmetric geometry and inhomogeneous surface charge density is not.^28^ Additionally, rectification ratios were found to be strongly correlated with elastomer choice (see Figure SI-B1), with much greater values observed.

To quantify the noise characteristics of a PEIN, we took the power spectrum at 100 kHz bandwidth for the bare nanopipette (before approaching the surface) at 5 mV, as well as PEINs of two resistances at 600 mV, which results in current values of the same order of magnitude. The resulting power spectral density (PSD) (Figure 1e) can be decomposed into its components *S*_*f*_ (*f* ^*−*1^), *S*_*w*_(*f* ^0^), *S*_*d*_(*f*) and *S*_*a*_(*f* ^2^), where *S*_*f*_, *S*_*w*_, *S*_*d*_ and *S*_*a*_ represent the PSD of 1*/f*, white, dielectric and amplifier noise respectively.^29^ At high frequencies, *f*, approaching the bandwidth, with noise dominated by the amplifier, the PSDs coincide. Similarly, the dielectric component, *S*_*d*_, is also constant with resistance as it is a function of total capacitance and the dielectric loss constant of the material which isolates the electrolyte. The defined peaks in the PSD are caused by the introduction of micromanipulators into the Faraday cage and could be removed by isolating them.

The white noise can be further decomposed into two sources: thermal and shot or Poisson noise:

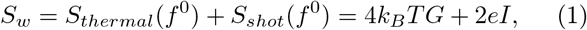

where *k*_*B*_ is the Boltzmann constant, *T* the temperature, *G* the conductance, *e* the elementary charge and *I* the current. Given the much larger aperture of the bare nanopipette, the resistance is significantly lower (1 MΩ; i.e., two orders of magnitude lower), and correspondingly the conductance higher. This is in quantitative agreement with the two-order-of-magnitude difference in the minima of the noise PSD floors of a 300 MΩ PEIN and bare nanopipette (Figure 1e).

The noise PSDs differ further in the low frequency regime where 1*/f* noise dominates. Surface charge fluctuations have been shown to cause 1*/f* noise, with material choice impacting these fluctuations.^29^ In particular, nanopore hydrophobicity increases 1*/f* noise,^30^ contributing to the low-frequency increase in a PEIN. Another contributor to 1*/f* noise is mechanical fluctuations, with more mechanically flexible nanopores exhibiting increased noise at low frequencies.^31^ Despite the use of an optical table, vibrations are still transmitted to PEINs, contributing to increased 1*/f* noise, as well as modulating the measured current with vibrational frequencies. This modulation is primarily seismic and can be seen in the PSD plot as a broad peak centred on ~ 19 Hz. As discussed earlier, this can be explained as the nanopipette and PDMS oscillating out of phase. At higher resistances, where the pipette and PDMS are more closely mechanically coupled (pentagon to hexagon in Figure 1b & c), 1*/f* noise and the broad peak are reduced, consistent with this logic.

### DNA Translocation

Having established the dynamics of nanopore formation, we studied the transport in further detail. As DNA is negatively charged, translocations through the nanopore are established simply by applying a potential, electrophoretically driving molecules through. Example traces for 1 kilo-base pair (kbp) dsDNA translocating through a PEIN (pipette to bath) in 2 M KCl 1 × TE (pH 9) at various resistances are shown in Figure 2b. Ostensibly, the measured traces are canonical for 1 kbp dsDNA in a glass nanopipette; however, their molecular dwell times exceed typical values by two orders of magnitude.^32^ These low speeds may arise from increased interaction between the DNA polymer and the hydrophobic PDMS surface: either directly due to the hydrophobicity of bases, or indirectly due to partial ordering of water molecules near the PDMS, increasing molecular friction as the polymer moves near the surface.^33^ However, this argument is not in accord with previous studies of the effect of hydrophobicity, where it was observed that hydrophilicity increases translocation times.^20^ Further discussion of the dynamics is withheld until a later section, where the increased dwell time with resistance is quantified.

**Figure 2:**
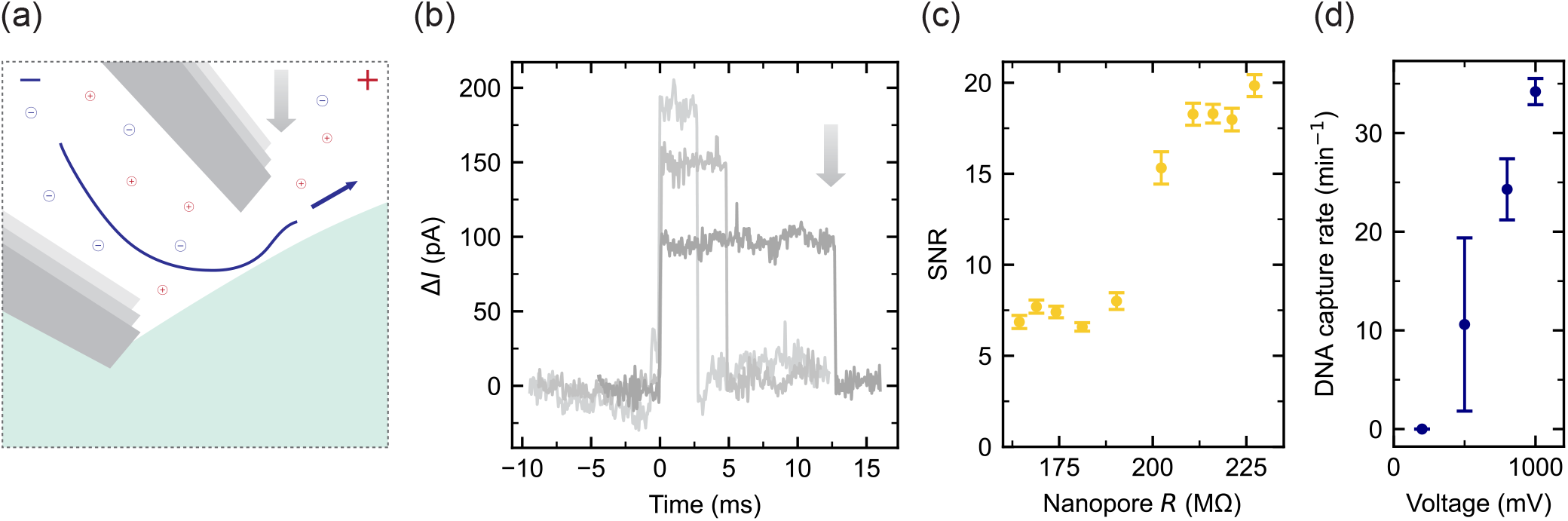
(a) Schematic of DNA translocating a pipette-elastomer interfacial nanopore (PEIN). (b) Representative current traces for 1 kbp double-stranded (ds)DNA translocating the PEIN from pipette to bath in 2 M KCl 1 × TE (pH 9). Darker shading corresponds to an increased resistance pore. (c) Signal-to-noise ratio (SNR): ratio of the average change in current during 1 kbp dsDNA translocation to the standard deviation of baseline before event, as a function of the resistance at which the translocation occurs at (at 800 mV). (d) Capture rate of 1 kbp dsDNA as a function of applied voltage.

Another unique characteristic of an interfacial nanopore is that these events are conductive in high salt; that is, DNA translocations cause events of increased current magnitude. This is contrary to the accepted rhetoric that in high salt, the presence of a DNA molecule in the nanopore reduces ion mobility in its vicinity, causing a resistive event.^34^ Although observed previously, no convincing argument was presented for conductive events in interfacial nanopores.^21–23^ Similar to our proposed explanation for low DNA velocities, ion mobility may already be reduced near the PDMS surface, and therefore the DNA-induced mobility reduction is less significant. The charge carrier density is higher at all salt concentrations when DNA translocates,^34^ and therefore a smaller relative reduction, or even increase, in ion mobility could lead to conductive events. Moreover, partial ordering of water molecules near PDMS could also lead to poorer hydration of charge carriers, resulting in a lower local concentration prior to DNA’s entry into the pore, also consistent with the slight ICR. More generally, DNA may be transiently wetting the pore, increasing current while it is present. This rationale is further consistent with the observed variability in relative change in current (|Δ*I/I*_0_|) between experiments: although typically for dsDNA |Δ*I/I*_0_| = 0.1, this can rise as high as 0.4 in some repeats. This effect may be the result of variability in the wetting of PDMS, enhancing the conductivity of the event. Within one repeat, however, relative current scales with resistance (*R*). Similarly, the signal-to-noise ratio (SNR), quantified as a function of resistance in Figure 2c, can vary significantly between experiments. Nonetheless, we observe an improvement in SNR with increasing resistance, as is expected from our analysis of the PSD.

We characterised the capture rate for 1 nM DNA simply by averaging the number of events per minute (Figure 2d). The rate exhibits the expected dependence on voltage, and agrees quantitatively with translocation frequencies for glass nanopipettes.^35^ However, while a lack of significant barrier to translocation was observed for nanopipettes, the opposite is observed for PEINs. If we again postulate that the interior of the nanopore is poorly hydrated due to the hydrophobic PDMS, there would then be an associated energy barrier for DNA to move into the interior. Furthermore, although there is no concentration imbalance between baths, a lower local concentration of charge carriers would increase the Debye layer thickness, potentially introducing an electro-osmotic flow (EOF) due to a preferred ion region near the negatively charged glass. This could then also contribute to reduced DNA velocities, as the EOF would oppose the electrophoretic driving of DNA, potentially preventing translocation at voltages below 200 mV.

### Molecular Transport

As discussed, a seminal property of PEINs is the dynamic aperture sizing. We therefore measured the dwell times for 1 kbp dsDNA and the resistance at which the translocation occurred (taken from the baseline current before the event). Sweeping across a range of ~ 100 MΩ, we could collect multiple thousand such pairings, with the results shown in Figure 3a (see Experimental Section for how the population was binned).

**Figure 3:**
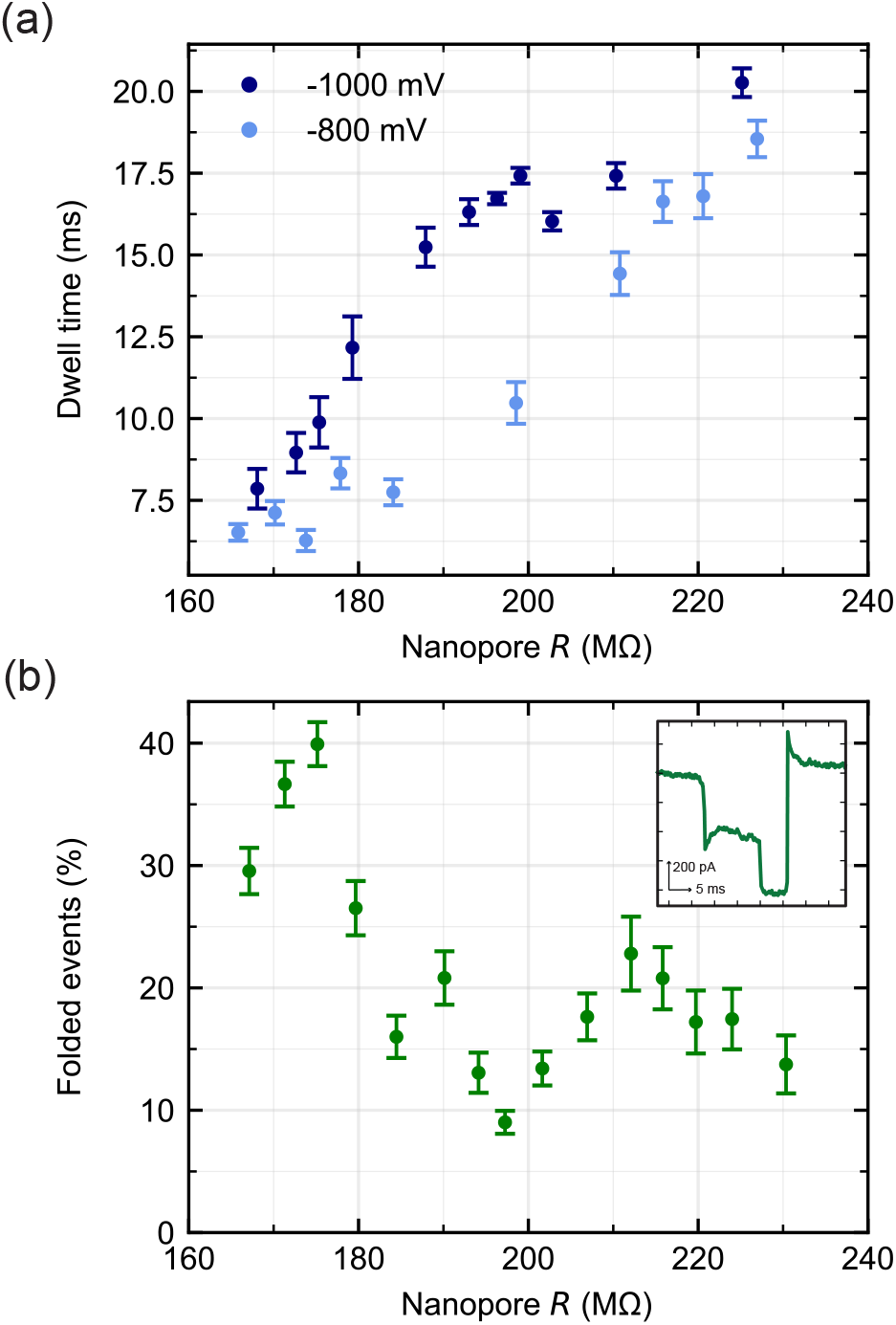
(a) Average dwell times for 1 kbp DNA for binned groups of events with resistance at which the DNA translocates and for two voltages. Each data point is derived from a minimum of 140 events. (b) Folding percentage of DNA events with resistance, combined for all voltages. Inset: representative trace for a folded event.

Within this ~ 100 MΩ window, we see the dwell time more than double. It should be noted that we did not filter out folded or knotted events. Such polymer conformations are characteristic of dsDNA translocating a solid-state nanopore, and we quantify the propensity of folding in PEINs in Figure 3b, with an example of a folded event shown in the inset. This quantification relies on the presence of a peak in standard deviation within a translocation event, and therefore the plateau in folded fraction at 10-20% corresponds to peaks in the event (see Figure SI-C1 for example traces). While these could be random noise rather than true polymer folding or knotting, particularly given the expected knotted fraction for 1 kbp is close to zero,^36^ increased rates of this kind of event have been observed for DNA translocating from a nanopipette to a bath, and so may correspond to a physical effect.^32^ Furthermore, we note the non-linear I-V curves prevent the use of resistance as a constant proxy for aperture size when comparing between voltages, as a 200 MΩ aperture measured at 1000 mV will be smaller than one measured at 800 mV, hence why 1000 mV appears to have longer dwell times in Figure 3a.

The observed increase in dwell time is consistent with correspondingly smaller nanopores,^19^ which reduce the folding tendency and increase the drag on the DNA. Above ~ 180 MΩ, there is no significant change in folding rate (Spearman’s *ρ* = 0.010, *p* = 0.490, *n* = 4558), and therefore only the latter effect contributes to the continued trend. Frictional drag in the pore should scale as:

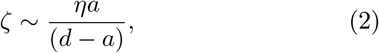

where *a* = 2.2 nm is the hydrodynamic cross section of DNA, *η* is the solvent viscosity and *d* the pore diameter.^19^ If we then approximate a PEIN as a cone with circular aperture whose resistance scales as *R ~* 1*/*(*πσd* tan *α*), where *σ* is the solution conductivity and *α* the conical angle,^26^ the drag coefficient scales as:

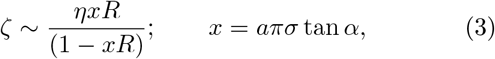

using a conductivity of 19 Sm^*−*1^ for 2 M KCl,^37^ a viscosity of *η* = 887.0 *µ*Pa.s,^38^ and *α≈* 12°, we expect, in this heavily simplified model, only a ~ 28% increase in dwell time in the range where events are linear ([180, 230] MΩ and using Stokes’ Law). Ignoring the simplicity of the employed model, there is clearly a stronger dependence of dwell time on resistance not captured by a frictional model alone.

Above ~ 230 MΩ, no events were observed, suggesting the aperture was constricted such that the minor radius of the aperture ellipse is less than the hydrodynamic radius of dsDNA. We use this trait in a later section to size the aperture’s minor axis using molecules of known radii. At lower resistances, the dwell times for different voltages converge as the non-linearity of the I-V curves reduces with resistance, becoming Ohmic when the nanopipette is far from the surface.

To further investigate molecular transport, we used a DNA origami ‘ruler’ introduced by Bell *et al*.,^32,39^ to study the velocity profile of a DNA polymer in a PEIN; that is, how the velocity of the polymer changes during its translocation of the pore. This was achieved by the self-assembly of six equally spaced groups of DNA ‘dumbbells’ onto a scaffold, forming the ruler (Figure 4a). The additional protrusions along the otherwise double-stranded DNA produce secondary current peaks when translocating a nanopore (Figure 4b; filtered down to 1 kHz for clarity), and given the prior knowledge of their distance in space, the measured distance in time between successive peaks yields the velocity profile of the DNA.

**Figure 4:**
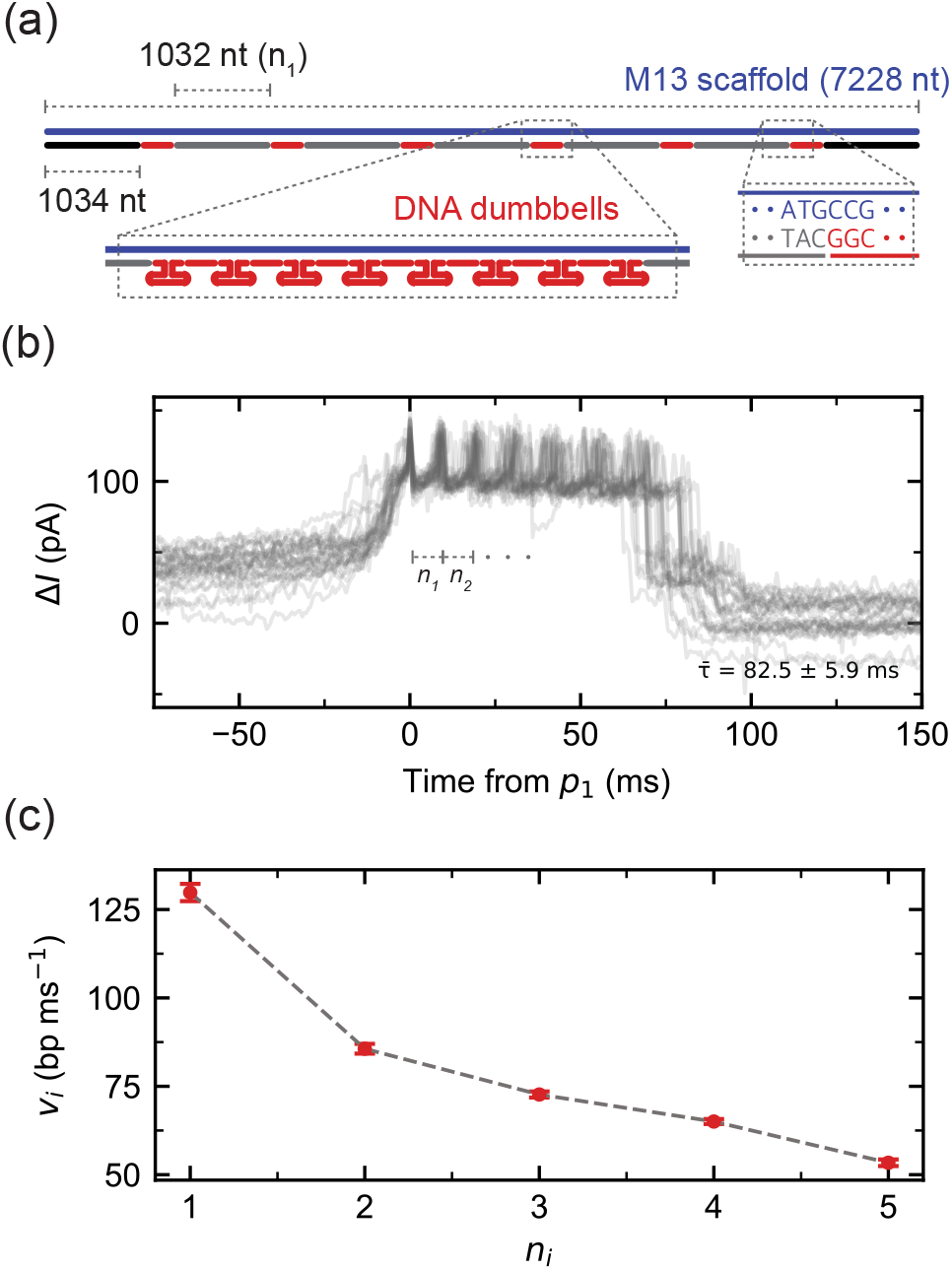
(a) Schematic of DNA origami structure: a 7228 nucleotide (nt) linearised M13 carrier with six groups of eight DNA dumbbells hybridised to it, with the groups separated by 1032 nt. (b) 30 overlaid raw example traces, filtered at 1 kHz, demonstrating the additional current peaks in the trace, aligned to the first peak. (c) Mean DNA velocity as translocation progresses calculated from the time between successive peaks, measured in base pairs (bp) per millisecond.

After assembly of the DNA rulers, they were diluted into the nanopipette and a negative voltage bias applied (1000 mV), with the resulting events recorded. We then averaged the velocities between peaks for many such events, with the results shown in Figure 4c. As discussed, velocity is resistance dependent, and thus in this case the resistance was maintained at a constant 200 MΩ. However, the continuous nature of the measurement produces a distribution of resistances around this target (standard deviation, *σ* = 7 MΩ); drift and stochastic changes in pore size mean each polymer translocates a different sized pore. Here we simply ignore this effect and average interpeak speeds, but the consequence is visible in Figure 4b where peaks become misaligned and six peaks are not clearly visible. Nonetheless, a clear trend in polymer velocities is observable: a non-linear reduction in speed.

While previous studies with symmetric pores observed the contrary, attributing a speeding up at the end of the translocation to the unfolding of the DNA,^40^ Bell *et al*. demonstrated that nanopore geometry can affect the velocity profile.^32^ Nevertheless, several differences with this work and previous studies should be noted. One such consideration is whether the DNA dwell time is less than the timescale of its conformational fluctuations, the longest wavelength of which is quantified using the Zimm relaxation time *τ*_*z*_, and thus does not equilibrate before the translocation is complete. DNA in concentrated monovalent salt solutions remains soluble and hydrated, and we therefore treat a 2 M KCl buffer as a good solvent in which the Zimm relaxation time is given by the expression:

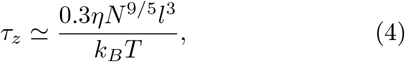

where *N* is the number of segments in the polymer and *l* the Kuhn length.^41^ Using a Kuhn length of 60 nm,^42^ the relaxation time for a 7.2 kbp DNA is 11 ms at room temperature. The relaxation time is then shorter than the translocation time, *τ*_*trans*_, while previous literature worked in the *τ*_*z*_ *> τ*_*trans*_ regime.

Bell *et al*. argued that only the straight portions of the polymer contribute to the net hydrodynamic force, as contributions from the fluctuating parts cancel in the vector sum of net force.^32^ Therefore, in their analogous case of DNA translocating from nanopipette to bath, the DNA is under tension and thus as translocation progresses, the length in the pore contributing to drag reduces, resulting in an increasing velocity. Whereas, in a PEIN, the polymer remains in equilibrium, and we therefore postulate that only the section translocating the pore is linear and contributing to the net drag. This would also explain why there is no acceleration as the tail of the polymer unravels: the unfolding of the DNA is not limiting its translocation. Moreover, the bath side of the pipette in a PEIN is relatively confined by the PDMS and thus the DNA, after translocating, buckles under compression. Increased crowding of DNA on the trans side could then inhibit further translocation, contributing to the observed decrease in velocity.

Regardless, this is further evidence that a hydrodynamic model of translocation is not sufficient to encapsulate the observed translocation dynamics, and that more complex molecule-surface interactions are present. We postulate that these interactions could, in signal-dimensionality terms, beneficially modify molecular transport in other biopolymers.

### Molecular Size: Sensitivity and Selectivity

Having established the influence of aperture size on molecular transport, we next sought to determine the extent to which the PEIN is sensitive to molecular dimensions. Specifically, we examined both the dependence of dwell time on polymer contour length and the extent to which the nanopore could discriminate analytes on the basis of their cross-sectional size.

We studied the scaling of dwell time with DNA length, *n*, (kbp) using dsDNA fragments between 0.5 and 5 kbp (Figure 5a) at 800 mV. To remove the dependence on resistance and folding propensity, the dwell times and errors for each length were obtained by averaging within a *±*1 MΩ window around 190 MΩ, yielding a superlinear scaling (*t ~ n*^*α*^, *α* = 1.20 *±* 0.07, *R*^2^ = 0.99). Comparable superlinear exponents have been reported, ranging from 1.10 up to 1.50,^32,43,44^ maintaining the persistence of *α* across nanopore geometries.

**Figure 5:**
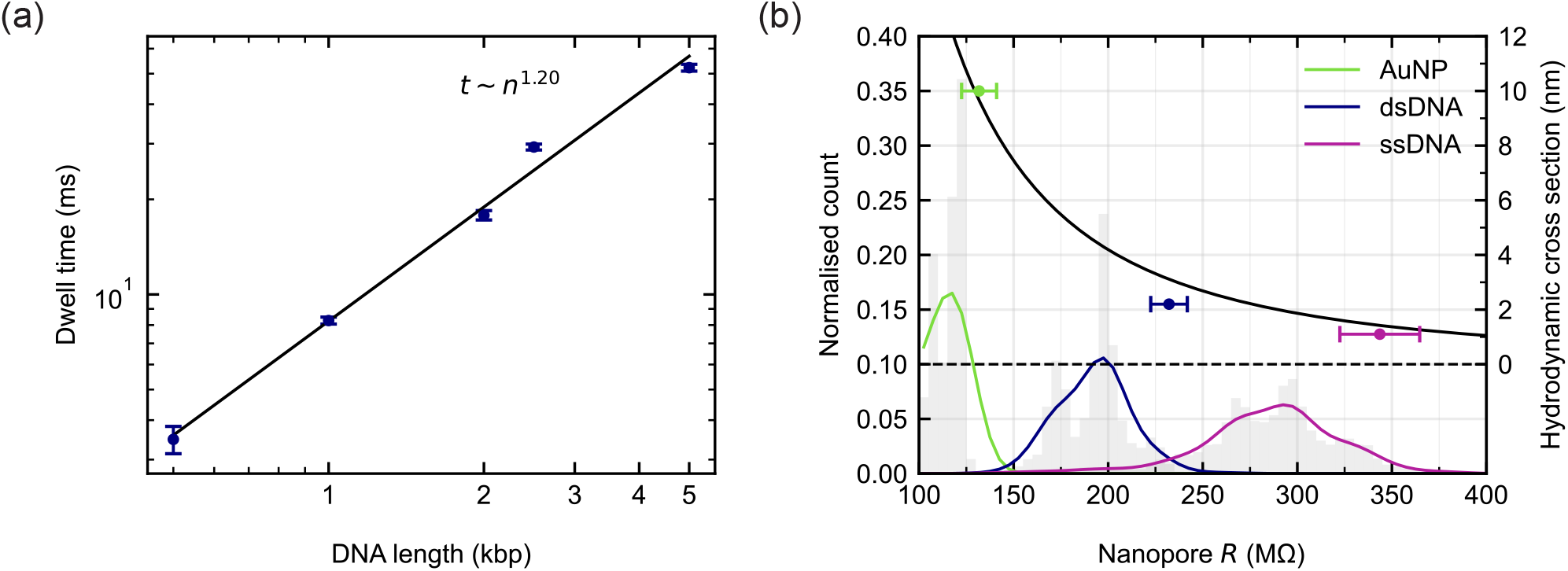
(a) Dwell time as a function of DNA length (kbp) of double-stranded (ds)DNA, calculated by averaging a resistance-dwell time plot at 190 MΩ. (b) Histograms of molecular populations by type, normalised by number of events of that type (grey). Smoothed curves are fitted to these histograms. The resistance at half maximum with the corresponding hydrodynamic cross section of the molecular type is shown in the overlaid scatter plot, with a least squares regression 1*/R*^2^ curve fitted (*R*^2^ = 0.98).

We demonstrate the ability to selectively gate the pore by molecular size by sweeping the resistance and recording the range over which events are observed. This was done for 10 nm gold nanoparticles and ss- and ds-DNA (Figure 5b). Resistance values were grouped into bins and normalised by the total number of events for each sample. The distributions were smoothed using a Gaussian kernel with a standard deviation of two bins. The resistance at the right-hand half-width half-maximum (HWHM) of each distribution was then determined by linear interpolation. Hydrodynamic cross sections of 1.1 nm, 2.2 nm and 10 nm for the ssDNA, dsDNA and nanoparticles respectively were plotted at the corresponding resistance, with horizontal error bars indicating the right-hand HWHM. If we arbitrarily assume the molecule size at this HWHM cut-off is equal to the aperture size (the molecule does not ‘fit’ through at higher resistances), we have an empirically derived calibration curve which can then be used to relate resistance to aperture size. Unsurprisingly, the relation is non-linear, with our naive scaling used in Equation 3 suggesting a 1*/R* relation, but this has a poor fit (*R*^2^ = 0.72). Instead we fit a polynomial, of order two, which is overlaid in Figure 5b (*R*^2^ = 0.98).

The distributions have cut-off points above which no events are observed, implying the aperture is too small to allow their translocation. The sharpness of the transition from allowed to not (average HWHM = 13.3 MΩ) implies the system is relatively rigid, and molecules cannot ‘squeeze’ through. The ssDNA distribution is broader (21.10 MΩ), perhaps corresponding to different foldings and hybridisations of the single-stranded polymer, with only more linear configurations permitted at higher resistances.

## Discussion

We demonstrated continuous control over molecular transport through a solid-state nanopore by adjusting the nanopore aperture *in situ*. DNA dwell times were systematically tuned over more than a twofold range due to the imposition of an increased degree of confinement experienced by translocating molecules, exceeding the predictions of simple frictional models, and DNA origami rulers revealed a progressive reduction in translocation velocity throughout passage of the pore. Together, these observations suggest that confinement within the PEIN modifies molecular transport beyond the effects expected from aperture size alone, highlighting the importance of surface interactions in determining translocation dynamics. Furthermore, we demonstrated that the PEIN can function as a dynamically adjustable molecular gate, selectively permitting or excluding molecules according to analyte size. Together, these results show that dynamic aperture sizing provides a direct means of controlling translocation dynamics beyond that achievable with conventional fixed geometry nanopores.

These capabilities are enabled by the pipette–elastomer interfacial nanopore, a dynamically reconfigurable nanopore developed in this work. A PEIN combines the simplicity and accessibility of glass nanopipette sensing with continuous, reversible aperture-size control while maintaining favourable electrical noise characteristics and reproducible operation, providing a single platform capable of interrogating a range of molecular sizes.

The ability to continuously tune nanopore confinement expands the experimental parameter space available for single-molecule measurements. Beyond improving control over translocation dynamics, size-selective gating offers a route towards isolating specific molecular populations and hints at the possibility of probing the conformational landscape of complex biopolymers, and perhaps inducing changes between conformational states. We anticipate that dynamically reconfigurable nanopores such as PEINs will provide a versatile platform for studying folded nucleic acids, proteins and other heterogeneous biopolymer systems, where the relationship between molecular conformation, transport dynamics and nanoscale confinement remains poorly understood.

## Experimental Section

### DNA Dumbbell Fabrication

The DNA scaffold, linearising stock oligonucleotides and dumbbell structures form the ‘DNA structure,’ developed in previous work by Bell *et al*. and was similarly used to characterise DNA velocity profiles.^32,39^ The 7228 nt scaffold was linearised from M13mp18 ss-DNA (7249 bases, N4040S, New England Biolabs) by hybridising a 39 nt oligonucleotide (5^*′*^-TCTAGAGGAT CCCCGGGTACCGAGCTCGAATTCGTAATC-3^*′*^, Integrated DNA Technologies) to the M13mp18 ssDNA by mixing 40 *µ*L M13mp18 ssDNA (250 ng/*µ*L), 8 *µ*L 10 ×Cutsmart buffer (New England Biolabs), 2 *µ*L the 39 nt oligonucleotide (100 *µ*M) and 28 *µ*L nuclease free water (NFW) before being heated to 65°C and linearly cooled down to 25°C in a thermocycler over 40 minutes. This partially hybridised scaffold was incubated with 1 *µ*L of the enzymes BamHI-HF and EcoRI-HF (each 100000 units/mL, New England Biolabs) at 37°C for one hour. A NucleoSpin Gel and PCR clean-up kit (MacheryNagel) was used to purify the cut scaffold, and its yield (DNA concentration) was quantified using a NanoDrop spectrophotometer.

The DNA structure was assembled by hybridising this 7228 nt single strand with short oligonucleotides in the ratio 1:5: 8 *µ*L linearised M13mp18 ssDNA (100 nM), 20 *µ*L oligonucleotide mixture (each oligo 200 nM, see Table SI-F1 for sequences), 4 *µ*L 100 mM MgCl_2_, 1.2 *µ*L 10 × TE (pH 8) and 6.8 *µ*L NFW. These were annealed at 70°C, followed by a linear cooling ramp to 25°C for 50 minutes, before being washed twice with 460 *µ*L 0.5 mM MgCl_2_,10 mM Tris-HCl (pH 8) buffer using an Amicon Ultra 100 kDa filter and centrifuged at 9000 × g for ten minutes at 4°C. The sample was then recovered by inverting the filter and centrifuging for one minute at 1000× g. The final yield was quantified using the NanoDrop. The DNA structures were stored at 4°C in 0.5 mM MgCl_2_, 10 mM Tris-HCl (pH 8) until measurement.

### Nanopore Fabrication

PDMS (Sylgard 184) was cast directly in Petri dishes with the curing agent to base ratio of 1:9 before degassing. The PDMS coated dishes were then cured at 80°C overnight. The PDMS was then further degassed and plasma treated (osupxygen and water), in order to prevent the nucleation and growth of microbubbles at the pipette-PDMS interface. However, while plasma treatment does increase the macroscopic hydrophilicity of the PDMS, its effects at the nanoscale could be variable, perhaps influencing molecular transport. Glass nanopipettes were fabricated from borosilicate capillaries (BF-150-86-10, World Precision Instruments) using a filament puller (P-97, Sutter Instruments). The pulling program was three-line: TIME = 250, P = 500, (1) HEAT = RAMP + 10, VEL = 28 (2) HEAT = RAMP + 10, VEL = 26, (3) HEAT = RAMP + 20, VEL = 28, PULL = 50.

### Nanopore Measurement

Analytes were diluted to 1 nM in a 2 M KCl 1× TE (pH 9) solution and pipetted into the nanopipettes. For Figure 2, Figure 3 and Figure 5a the analytes were dsDNA fragments (0.5-5 kbp, NoLimits DNA Fragments, Thermo Fisher) For the selectivity plot (Figure 5b) the analytes used were linearised M13mp18 ss-DNA (7249 bases, N4040S, New England Biolabs) and 10 nm gold nanoparticles (741967-25ML, Sigma-Aldrich). Aggregation of nanoparticles was limited by diluting into 2 M KCl only at the point of measurement. The ds-DNA here was self-assembled from the M13 ssDNA. The nanopipettes were mounted on an Ag/AgCl-electrode-containing pipette holder (EH-P170), which was in turn mounted on the amplifier headstage at 30°. An amplifier (Axopatch 200B, Molecular Devices) and hardware Bessel filter (3384, Krohn-Hite) were used to measure and filter (10 kHz) before the signal was digitised at 50 kHz (Molecular Devices 1440A, Molecular Devices). Translocation events were recorded using a custom C++ program developed to use a simple threshold to identify and record only events.

## Data Analysis

Data was cleaned of clogging (and folded) DNA translocation events using a custom Python program which filters for linear events using common event characteristics, leaving only readable DNA events (see SI Section E). Additional current peaks in the DNA ruler events, or for presence of folding, were found using a simple standard deviation peak finding algorithm. The relationship between pore resistance and dwell time/SNR was quantified using adaptive kernel-based binning along the resistance axis, with the initial number of bins estimated using the Freedman-Diaconis rule. Rather than constructing uniformly spaced histogram bins, a Gaussian kernel density estimate (KDE) of the resistance distribution was calculated and its cumulative distribution function was used to determine equal-probability bin boundaries. To prevent over-segmentation in densely populated regions, neighbouring bins were merged whenever the resulting bin width was less than 3 MΩ, while maintaining a minimum average occupancy of 140 events per bin. The mean SNR or dwell times and resistances of these bins, along with the standard error in the mean are the plotted points in Figure 2c and 3a. Because the measurement uncertainty of individual events was negligible compared with the observed interevent variability, the sample variance was taken to represent the underlying statistical dispersion.

## Acknowledgements

The authors thank the group of János Vörös for their hospitality, and Annina Stuber, Julian Hengsteler and members of the BMC group for their support and discussion. This research has received funding from the Swiss State Secretariat for Education, Research and Innovation (SERI) [MB23.00009], associated with the EU-funded project NANOMICS (ERC Starting Grant).

## Supporting information

The following files are available free of charge.

- SI.pdf: contains supporting figures covering pore formation (FEM modelling of PDMS deformation, DIC imaging of PEINs where translocations were observed and examples of failures, modelling of capillary bending), material effect (approach and IV curves for common elastomers), further example traces (when peaks in the event are observed for 1 kbp), reproducibility (repeated approach and retraction curves, decomposition of Figure 3a), data analysis (pipeline use to filter recorded DNA events from noise-related events) and the complete sequences of DNA used for self-assembly of the DNA origami.

