## Supporting Information for "Controlling Molecular Transport through Nanopores by Dynamic Aperture Sizing"

### A Pore Formation

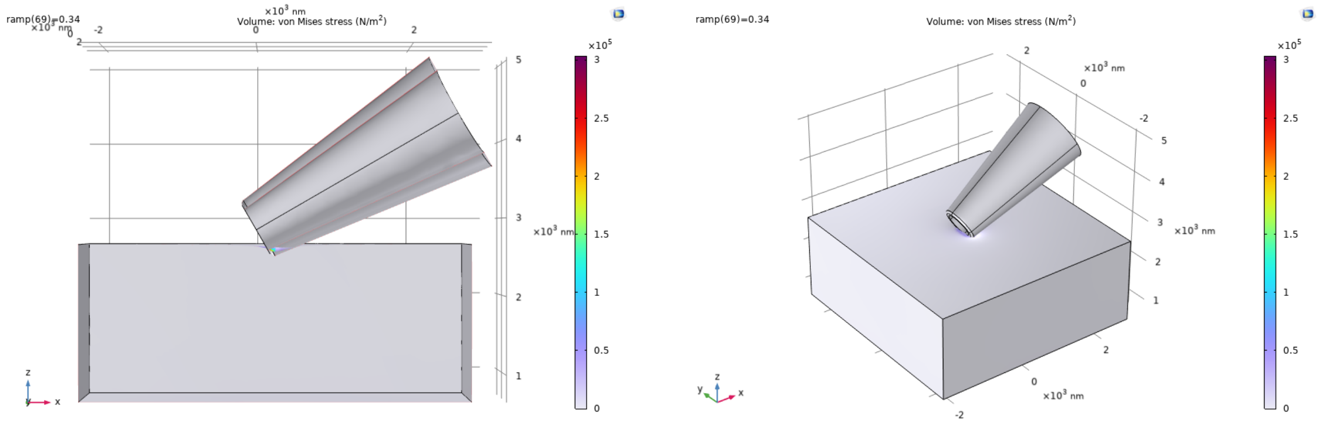


Figure A1: COMSOL finite-element simulations of a glass cone of similar dimension to glass nanopipettes used, showing the von Mises stress distribution and deformation of PDMS around the aperture and reducing the effective aperture size.

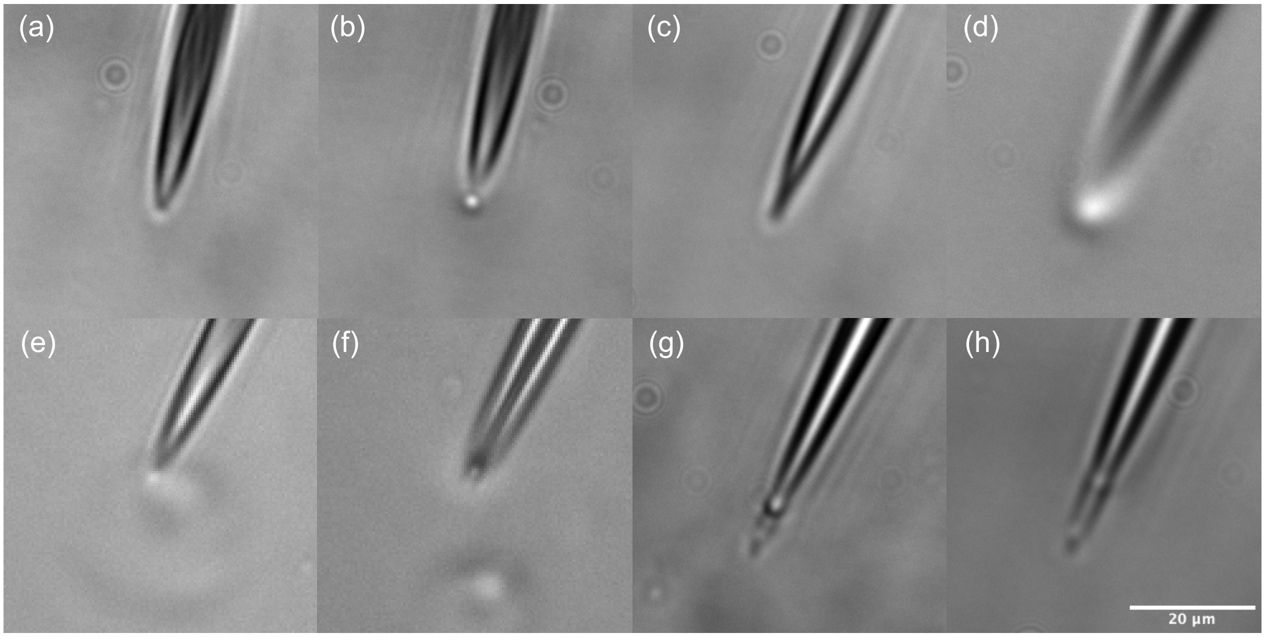


Figure A2: Differential interference contrast (DIC) images of glass nanopipettes pressing into a PDMS surface. (a)-(d) show images of the nanopore when events were observed, while (e)-(h) show images when no events were observed. (e) & (f) show a bubble at the tip, either captured from solution or directly nucleated at the PDMS-pipette interface. In (g) & (h), a shallower angle (30°) was used and show puncturing of the PDMS surface.

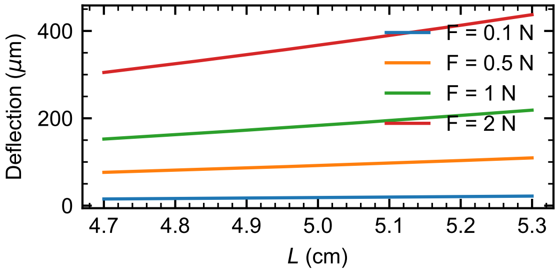


Figure A3: Deflection $\delta$ of glass capillary with force.

Modelling the capillary as a hollow cylindrical cantilever beam, the deflection due to a force applied to its end is $\delta = 4FL^3/3\pi E(r_o^4 - r_i^4)$, where $F$ is the applied force, $L$ the capillary length, $E$ the Young's Modulus of borosilicate glass and $r_o$ and $r_i$ are the inner and outer radii of the capillary. Figure A3 shows the deflection as a function of reasonable applied forces over the range of capillary lengths, $L$, that can arise from pipette pulling ($\pm 1$ mm).

### B Alternative Materials

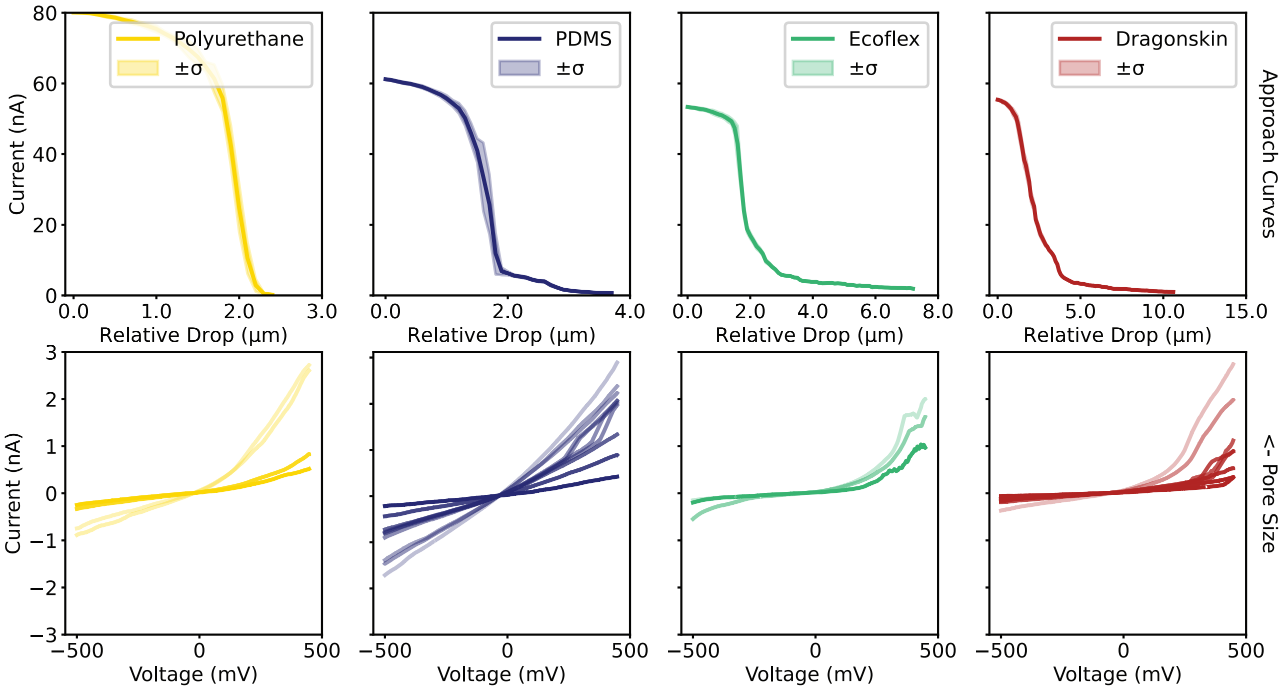


Figure B1: Exemplary approach curves (current with relative height drop from 98% initial current) for various materials in 2M KCl: Polyurethane (Tecothane TPU, TT95A, Lubrizol), PDMS (Sylgard 184), EcoFlex (Ecoflex 00-20, Smooth-On), Dragon skin (Dragon Skin 20, Smooth-On). Below are corresponding current-voltage curves for various pore sizes.

### C Example Traces

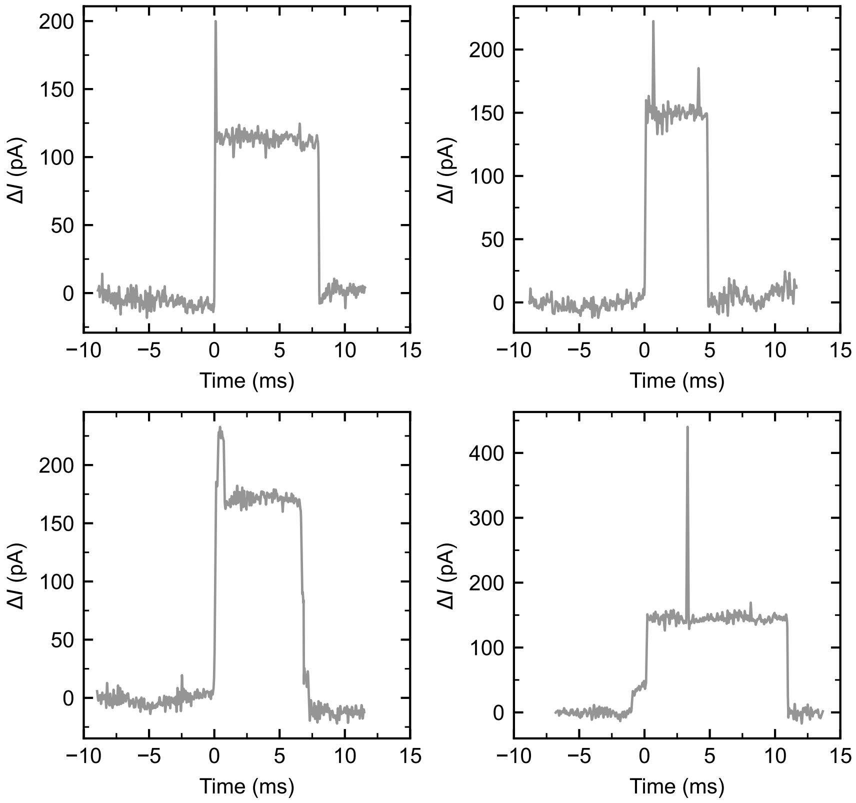


Figure C1: Example traces for 1 kbp double-stranded DNA (2 M KCl 1× TE (pH 9); 800 mV) where additional peaks were observed in the event.

### D Reproducibility

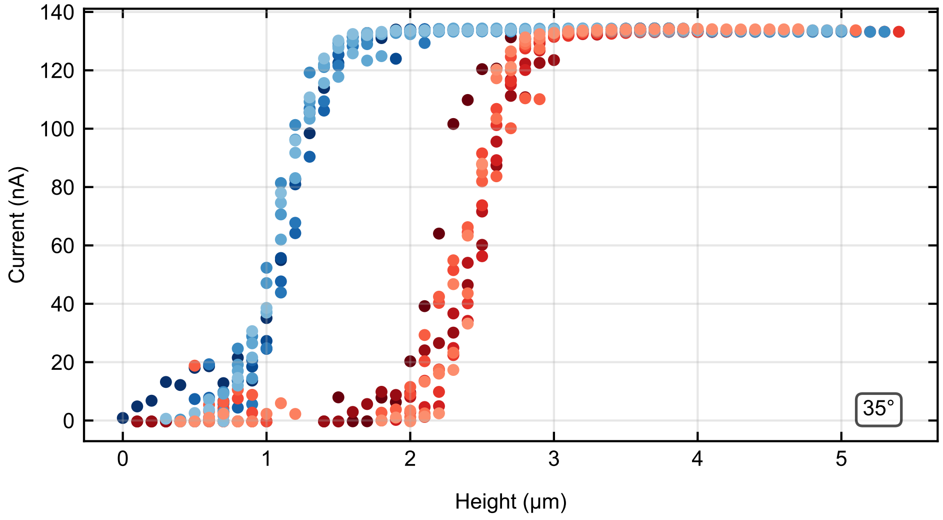


Figure D1: Ten consecutive approach (blue) and retractions (red).

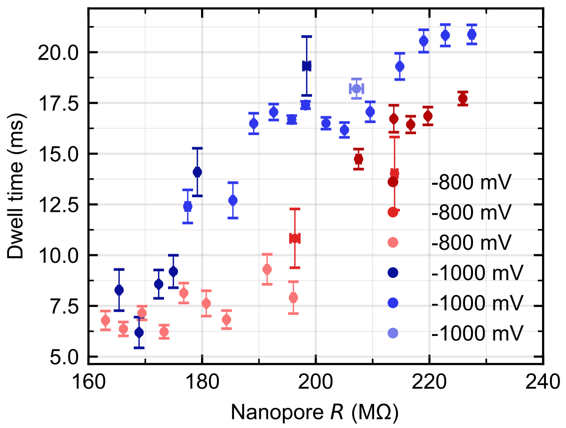


Figure D2: Shows Figure 3a decomposed into its contributing experiments.

### E Data Analysis

Translocation events were recorded using a simple thresholding algorithm, saving events where the current exceeded a user defined threshold (0.1 nA for DNA) measured from the current baseline (1 ms of averaged current data). A user defined region of context before and after this threshold exceeding region is appended to the recorded event (e.g. 10 ms before and after event). The resulting array of events is filtered to remove those events which correspond to noise rather than true translocations such as electrical noise, vibrations, clogging etc.. DNA translocations were extracted using common parameters (event charge, change in current etc.) by displaying the entire data set in two-parameter spaces, with translocations clustering in these spaces (Figure E1).

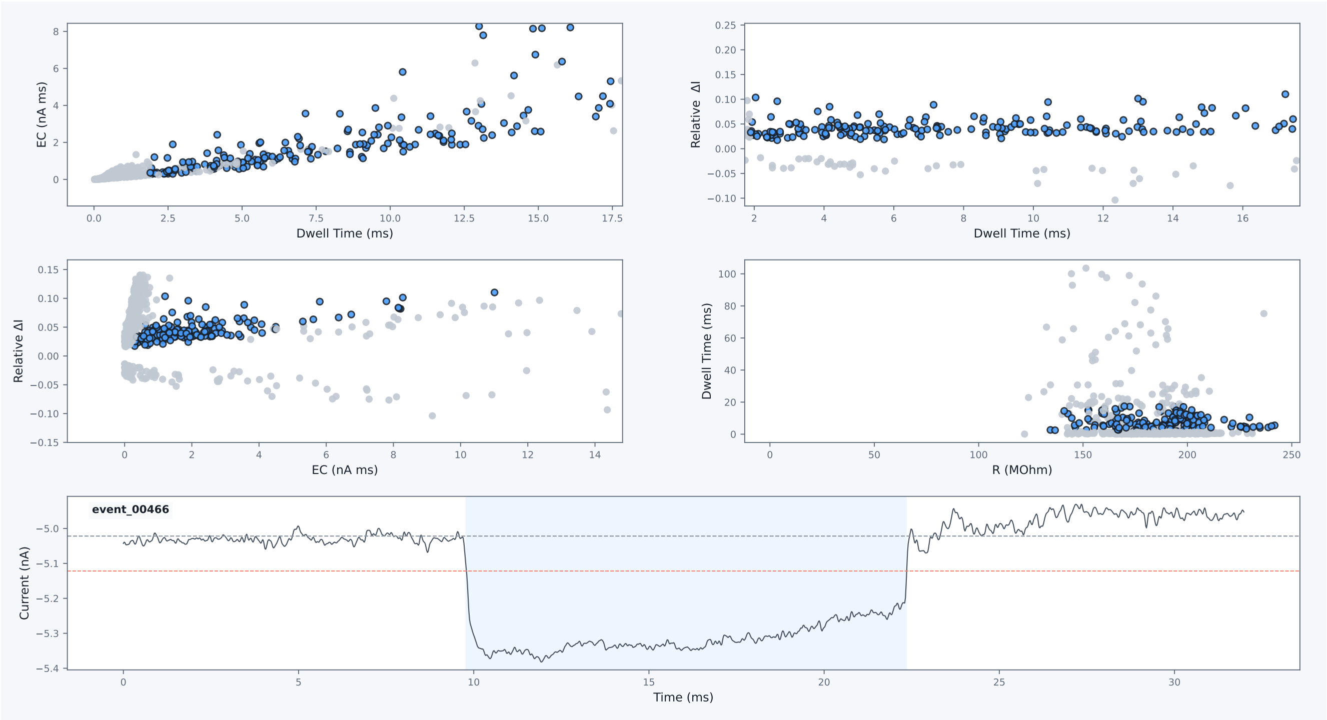


Figure E1: Screenshot of filtering UI, showing clustering of true translocation events in two-parameter spaces.

### F DNA Sequences

Table F1: DNA oligo sequences which were hybridised to the M13 carrier to form the DNA ruler.

| Position | Sequence (5' → 3') |
|---|---|
| 1 | TTTTCGTAATCATGGTCATAGCTGTTTCCTGTGTGAAATTGTTATC |
| 2 | CGCTCACAATTCCACACAACATACGAGCCGGAAGCATA |
| 3 | AAGTGTAAAGCCTGGGGTGCCTAATGAGTGAGCTAACT |
| 4 | CACATTAATTGCGTTGCGCTCACTGCCCGCTTTCCAGT |
| 5 | CGGGAAACCTGTCGTGCCAGCTGCATTAATGAATCGGC |
| 6 | CAACGCGCGGGGAGAGGCGGTTTGCGTATTGGGCGCCA |
| 7 | GGGTGGTTTTTCTTTTCACCAGTGAGACGGGCAACAGC |
| 8 | TGATTGCCCTTCACCGCCTGGCCCTGAGAGAGTTGCAG |
| 9 | CAAGCGGTCCACGCTGGTTTGCCCCAGCAGGCGAAAAT |
| 10 | CCTGTTTGATGGTGGTTCCGAAATCGGCAAAATCCCTT |
| 11 | ATAAATCAAAAGAATAGCCCGAGATAGGGTTGAGTGTT |
| 12 | GTTCCAGTTTGGAACAAGAGTCCACTATTAAAGAACGT |
| 13 | GGACTCCAACGTCAAAGGGCGAAAAACCGTCTATCAGG |
| 14 | GCGATGGCCCACTACGTGAACCATCACCCAAATCAAGT |
| 15 | TTTTTGGGGTCGAGGTGCCGTAAAGCACTAAATCGGAA |
| 16 | CCCTAAAGGGAGCCCCCGATTTAGAGCTTGACGGGGAA |
| 17 | AGCCGGCGAACGTGGCGAGAAAGGAAGGGAAGAAAGCG |
| 18 | AAAGGAGCGGGCGCTAGGGCGCTGGCAAGTGTAGCGGT |
| 19 | CACGCTGCGCGTAACCACCACACCCGCCGCGCTTAATG |
| 20 | CGCCGCTACAGGGCGCGTACTATGGTTGCTTTGACGAG |
| 21 | CACGTATAACGTGCTTTCCTCGTTAGAATCAGAGCGGG |
| 22 | AGCTAAACAGGAGGCCGATTAAAGGGATTTTAGACAGG |
| 23 | AACGGTACGCCAGAATCCTGAGAAGTGTTTTTATAATC |
| 24 | AGTGAGGCCACCGAGTAAAAGAGTCTGTCCATCACGCA |
| 25 | AATTAACCGTTGTAGCAATACTTCTTTGATTAGTAATA |
| 26 | ACATCACTTGTCCTCTTTTGAGGAACAAGTTTTCTTGTCCTGAGTAGA |
| 27 | AGAACTCAAATCCTCTTTTGAGGAACAAGTTTTCTTGTCTATCGGCCT |
| 28 | TGCTGGTAATTCCTCTTTTGAGGAACAAGTTTTCTTGTATCCAGAACA |
| 29 | ATATTACCGCTCCTCTTTTGAGGAACAAGTTTTCTTGTCAGCCATTGC |
| 30 | AACAGGAAAATCCTCTTTTGAGGAACAAGTTTTCTTGTACGCTCATGG |
| 31 | AAATACCTACTCCTCTTTTGAGGAACAAGTTTTCTTGTATTTTGACGC |
| 32 | TCAATCGTCTTCCTCTTTTGAGGAACAAGTTTTCTTGTGAAATGGATT |
| 33 | ATTTACATTGTCCTCTTTTGAGGAACAAGTTTTCTTGTGCAGATTCAC |
| 34 | CAGTCACACGACCAGTAATAAAAGGGACAT |
| 35 | TCTGGCCAACAGAGATAGAACCCTTCTGACCTGAAAGC |
| 36 | GTAAGAATACGTGGCACAGACAATATTTTTGAATGGCT |
| 37 | ATTAGTCTTTAATGCGCGAACTGATAGCCCTAAAACAT |
| 38 | CGCCATTAAAAATACCGAACGAACCACCAGCAGAAGAT |
| 39 | AAAACAGAGGTGAGGCGGTCAGTATTAACACCGCCTGC |
| 40 | AACAGTGCCACGCTGAGAGCCAGCAGCAAATGAAAAAT |
| 41 | CTAAAGCATCACCTTGCTGAACCTCAAATATCAAACCC |
| 42 | TCAATCAATATCTGGTCAGTTGGCAAATCAACAGTTGA |
| 43 | AAGGAATTGAGGAAGGTTATCTAAAATATCTTTAGGAG |
| 44 | CACTAACAACTAATAGATTAGAGCCGTCAATAGATAAT |
| 45 | ACATTTGAGGATTTAGAAGTATTAGACTTTACAAACAA |
| 46 | TTCGACAACTCGTATTAAATCCTTTGCCCGAACGTTAT |
| 47 | TAATTTTAAAAGTTTGAGTAACATTATCATTTTGCGGA |
| 48 | ACAAAGAAACCACCAGAAGGAGCGGAATTATCATCATA |
| 49 | TTCCTGATTATCAGATGATGGCAATTCATCAATATAAT |
| 50 | CCTGATTGTTTGGATTATACTTCTGAATAATGGAAGGG |

Continued on next page

Table F1 – continued from previous page

| Position | Sequence (5' → 3') |
|---|---|
| 51 | TTAGAACCTACCATATCAAAATTATTTGCACGTAAAAC |
| 52 | AGAAATAAAGAAATTGCGTAGATTTTCAGGTTTAACGT |
| 53 | CAGATGAATATACAGTAACAGTACCTTTTACATCGGGA |
| 54 | GAAACAATAACGGATTCGCCTGATTGCTTTGAATACCA |
| 55 | AGTTACAAAATCGCGCAGAGGCGAATTATTCATTTCAA |
| 56 | TTACCTGAGCAAAAGAAGATGATGAAACAAACATCAAGAAAACA |
| 57 | AAATTAATTATCCTCTTTTGAGGAACAAGTTTTCTTGTCATTTAACAA |
| 58 | TTTCATTTGATCCTCTTTTGAGGAACAAGTTTTCTTGTATTACCTTTT |
| 59 | TTAATGGAAATCCTCTTTTGAGGAACAAGTTTTCTTGTCAGTACATAA |
| 60 | ATCAATATATTCCTCTTTTGAGGAACAAGTTTTCTTGTGTGAGTGAAT |
| 61 | AACCTTGCTTTCCTCTTTTGAGGAACAAGTTTTCTTGTCTGTAAATCG |
| 62 | TCGCTATTAATCCTCTTTTGAGGAACAAGTTTTCTTGTTTAATTTTCC |
| 63 | CTTAGAATCCTCCTCTTTTGAGGAACAAGTTTTCTTGTTTGAAAACAT |
| 64 | AGCGATAGCTTCCTCTTTTGAGGAACAAGTTTTCTTGTTAGATTAAGA |
| 65 | CGCTGAGAAGAGTCAATAGTGAAT |
| 66 | TTATCAAAATCATAGGTCTGAGAGACTACCTTTTTAAC |
| 67 | CTCCGGCTTAGGTTGGGTTATATAACTATATGTAAATG |
| 68 | CTGATGCAAATCCAATCGCAAGACAAAGAACGCGAGAA |
| 69 | AACTTTTTCAAATATATTTTAGTTAATTTCATCTTCTG |
| 70 | ACCTAAATTTAATGGTTTGAAATACCGACCGTGTGATA |
| 71 | AATAAGGCGTTAAATAAGAATAAACACCGGAATCATAA |
| 72 | TTACTAGAAAAAGCCTGTTTAGTATCATATGCGTTATA |
| 73 | CAAATTCTTACCAGTATAAAGCCAACGCTCAACAGTAG |
| 74 | GGCTTAATTGAGAATCGCCATATTTAACAACGCCAACA |
| 75 | TGTAATTTAGGCAGAGGCATTTTCGAGCCAGTAATAAG |
| 76 | AGAATATAAAGTACCGACAAAAGGTAAAGTAATTCTGT |
| 77 | CCAGACGACGACAATAAACAACATGTTCAGCTAATGCA |
| 78 | GAACGCGCCTGTTTATCAACAATAGATAAGTCCTGAAC |
| 79 | AAGAAAAATAATATCCCATCCTAATTTACGAGCATGTA |
| 80 | GAAACCAATCAATAATCGGCTGTCTTTCCTTATCATTC |
| 81 | CAAGAACGGGTATTAAACCAAGTACCGCACTCATCGAG |
| 82 | AACAAGCAAGCCGTTTTTATTTTCATCGTAGGAATCAT |
| 83 | TACCGCGCCCAATAGCAAGCAAATCAGATATAGAAGGC |
| 84 | TTATCCGGTATTCTAAGAACGCGAGGCGTTTTAGCGAA |
| 85 | CCTCCCGACTTGCGGGAGGTTTTGAAGCCTTAAATCAA |
| 86 | GATTAGTTGCTATTTTGCACCCAGCTACAATTTTATCC |
| 87 | TGAATCTTACCAACGCTAACGAGCGTCTTTCCAGAGCCTAATTTGCCAGT |
| 88 | TACAAAATAATCCTCTTTTGAGGAACAAGTTTTCTTGTACAGCCATAT |
| 89 | TATTTATCCCTCCTCTTTTGAGGAACAAGTTTTCTTGTAATCCAAATA |
| 90 | AGAAACGATTTCCTCTTTTGAGGAACAAGTTTTCTTGTTTTTGTTTAA |
| 91 | CGTCAAAAATTCCTCTTTTGAGGAACAAGTTTTCTTGTGAAAATAGCA |
| 92 | GCCTTTACAGTCCTCTTTTGAGGAACAAGTTTTCTTGTAGAGAATAAC |
| 93 | ATAAAAACAGTCCTCTTTTGAGGAACAAGTTTTCTTGTGGAAGCGCAT |
| 94 | TAGACGGGAGTCCTCTTTTGAGGAACAAGTTTTCTTGTAATTAACTGA |
| 95 | ACACCCTGAATCCTCTTTTGAGGAACAAGTTTTCTTGTCAAAGTCAGA |
| 96 | GGGTAATTGAGCGCTAATATCAGAGAGATAACCCACAAGAATTGAGTTAAGCCCAA |
| 97 | TAATAAGAGCAAGAAACAATGAAATAGCAATAGCTATC |
| 98 | TTACCGAAGCCCTTTTTAAGAAAAGTAAGCAGATAGCC |
| 99 | GAACAAAGTTACCAGAAGGAAACCGAGGAAACGCAATA |
| 100 | ATAACGGAATACCCAAAAGAACTGGCATGATTAAGACT |
| 101 | CCTTATTACGCAGTATGTTAGCAAACGTAGAAAATACA |
| 102 | TACATAAAGGTGGCAACATATAAAAGAAACGCAAAGAC |

Continued on next page

| Position | Sequence (5' → 3') |
|---|---|
| 103 | ACCACGGAATAAGTTTATTTTGTCACAATCAATAGAAA |
| 104 | ATTCATATGGTTTACCAGCGCCAAAGACAAAAGGGCGA |
| 105 | CATTCAACCGATTGAGGGAGGGAAGGTAAATATTGACG |
| 106 | GAAATTATTCATTAAAGGTGAATTATCACCGTCACCGA |
| 107 | CTTGAGCCATTTGGGAATTAGAGCCAGCAAAATCACCA |
| 108 | GTAGCACCATTACCATTAGCAAGGCCGGAAACGTCACC |
| 109 | AATGAAACCATCGATAGCAGCACCGTAATCAGTAGCGA |
| 110 | CAGAATCAAGTTTGCCTTTAGCGTCAGACTGTAGCGCG |
| 111 | TTTTCATCGGCATTTTCGGTCATAGCCCCCTTATTAGC |
| 112 | GTTTGCCATCTTTTCATAATCAAAATCACCGGAACCAG |
| 113 | AGCCACCACCGGAACCGCCTCCCTCAGAGCCGCCACCC |
| 114 | TCAGAACCGCCACCCTCAGAGCCACCACCCTCAGAGCC |
| 115 | GCCACCAGAACCACCACCAGAGCCGCCGCCAGCATTGA |
| 116 | CAGGAGGTTGAGGCAGGTCAGACGATTGGCCTTGATAT |
| 117 | TCACAAACAAATAAATCCTCATTAAAGCCAGAATGGAA |
| 118 | AGCGCAGTCTCTGAATTTACCGTTCCAGTAAGCGTCAT |
| 119 | ACATGGCTTTTGATGATACAGGAGTGTACTGGTAATAA |
| 120 | GTTTTAACGGGGTCAGTGCCTTGAGTAACAGTGCCCGT |
| 121 | ATAAACAGTTAATGCCCCCTGCCTATTTCGGAACCTAT |
| 122 | TATTCTGAAACATGAAAGTATTAAGAGGCTGAGACTCC |
| 123 | TCAAGAGAAGGATTAGGATTAGCGGGGTTTTGCTCAGT |
| 124 | ACCAGGCGGATAAGTGCCGTCGAGAGGGTTGATATAAG |
| 125 | TATAGCCCGGAATAGGTGTATCACCGTACTCAGGAGGT |
| 126 | TTAGTACCGCCACCCTCAGAACCGCCACCCTCAGAACC |
| 127 | TCACAAACAAATAAATCCTCATTAAAGCCAGAATGGAAAGCGCAGTCTCTGAATTT |
| 128 | ACCGTTCCAGTCCTCTTTTGAGGAACAAGTTTTCTTGTTAAGCGTCAT |
| 129 | ACATGGCTTTTCCTCTTTTGAGGAACAAGTTTTCTTGTTGATGATACA |
| 130 | GGAGTGTACTTCCTCTTTTGAGGAACAAGTTTTCTTGTGGTAATAAGT |
| 131 | TTTAACGGGGTCCTCTTTTGAGGAACAAGTTTTCTTGTTCAGTGCCTT |
| 132 | GAGTAACAGTTCCTCTTTTGAGGAACAAGTTTTCTTGTGCCCGTATAA |
| 133 | ACAGTTAATGTCCTCTTTTGAGGAACAAGTTTTCTTGTCCCCCTGCCT |
| 134 | ATTTCGGAACTCCTCTTTTGAGGAACAAGTTTTCTTGTCTATTATTCT |
| 135 | GAAACATGAATCCTCTTTTGAGGAACAAGTTTTCTTGTAGTATTAAGA |
| 136 | GGCTGAGACTCCTCAAGAGAAGGATTAGGATTAGCGGGGTTTTGCTCAGT |
| 137 | ATAATTTTTTCACGTTGAAAATCTCCAAAAAAAAAGGCT |
| 138 | CCAAAAGGAGCCTTTAATTGTATCGGTTTATCAGCTTG |
| 139 | CTTTCGAGGTGAATTTCTTAAACAGCTTGATACCGATA |
| 140 | GTTGCGCCGACAATGACAACAACCATCGCCCACGCATA |
| 141 | ACCGATATATTCGGTCGCTGAGGCTTGCAGGGAGTTAA |
| 142 | AGGCCGCTTTTGCGGGATCGTCACCCTCAGCAGCGAAA |
| 143 | GACAGCATCGGAACGAGGGTAGCAACGGCTACAGAGGC |
| 144 | TTTGAGGACTAAAGACTTTTTCATGAGGAAGTTTCCAT |
| 145 | TAAACGGGTAAAATACGTAATGCCACTACGAAGGCACC |
| 146 | AACCTAAAACGAAAGAGGCAAAAGAATACACTAAAACA |
| 147 | CTCATCTTTGACCCCCAGCGATTATACCAAGCGCGAAA |
| 148 | CAAAGTACAACGGAGATTTGTATC |
| 149 | ATCGCCTGATTCCTCTTTTGAGGAACAAGTTTTCTTGTAAATTGTGTC |
| 150 | GAAATCCGCGTCCTCTTTTGAGGAACAAGTTTTCTTGTACCTGCTCCA |
| 151 | TGTTACTTAGTCCTCTTTTGAGGAACAAGTTTTCTTGTCCGGAACGAG |
| 152 | GCGCAGACGGTCCTCTTTTGAGGAACAAGTTTTCTTGTTCAATCATAA |
| 153 | GGGAACCGAATCCTCTTTTGAGGAACAAGTTTTCTTGTCTGACCAACT |
| 154 | TTGAAAGAGGTCCTCTTTTGAGGAACAAGTTTTCTTGTACAGATGAAC |

Continued on next page

Table F1 – continued from previous page

| Position | Sequence (5' → 3') |
|---|---|
| 155 | GGTGTACAGATCCTCTTTTGAGGAACAAGTTTTCTTGTCCAGGCGCAT |
| 156 | AGGCTGGCTGTCCTCTTTTGAGGAACAAGTTTTCTTGTACCTTCATCA |
| 157 | AGAGTAATCTTGACAAGAACCGGATATTCATTACCCAAATCAAC |
| 158 | GACGAGAAACACCAGAACGAGTAGTAAATTGGGCTTGA |
| 159 | GATGGTTTAATTTCAACTTTAATCATTGTGAATTACCT |
| 160 | TATGCGATTTTAAGAACTGGCTCATTATACCAGTCAGG |
| 161 | ACGTTGGGAAGAAAAATCTACGTTAATAAAACGAACTA |
| 162 | ACGGAACAACATTATTACAGGTAGAAAGATTCATCAGT |
| 163 | TGAGATTTAGGAATACCACATTCAACTAATGCAGATAC |
| 164 | ATAACGCCAAAAGGAATTACGAGGCATAGTAAGAGCAA |
| 165 | CACTATCATAACCCTCGTTTACCAGACGACGATAAAAA |
| 166 | CCAAAATAGCGAGAGGCTTTTGCAAAAGAAGTTTTGCC |
| 167 | AGAGGGGGTAATAGTAAAATGTTTAGACTGGATAGCGT |
| 168 | CCAATACTGCGGAATCGTCATAAATATTCATTGAATCC |
| 169 | CCCTCAAATGCTTTAAACAGTTCAGAAAACGAGAATGA |
| 170 | CCATAAATCAAAAATCAGGTCTTTACCCTGACTATTAT |
| 171 | AGTCAGAAGCAAAGCGGATTGCATCAAAAAGATTAAGA |
| 172 | GGAAGCCCGAAAGACTTCAAATATCGCGTTTTAATTCG |
| 173 | AGCTTCAAAGCGAACCAGACCGGAAGCAAACTCCAACA |
| 174 | GGTCAGGATTAGAGAGTACCTTTAATTGCTCCTTTTGA |
| 175 | TAAGAGGTCATTTTTGCGGATGGCTTAGAGCTTAATTG |
| 176 | CTGAATATAATGCTGTAGCTCAACATGTTTTAAATATG |
| 177 | CAACTAAAGTACGGTGTCTGGAAGTTTCATTCCATATA |
| 178 | ACAGTTGATTCCCAATTCTGCGAACGAGTA |
| 179 | GATTTAGTTTTCCTCTTTTGAGGAACAAGTTTTCTTGTGACCATTAGA |
| 180 | TACATTTCGCTCCTCTTTTGAGGAACAAGTTTTCTTGTAAATGGTCAA |
| 181 | TAACCTGTTTTCCTCTTTTGAGGAACAAGTTTTCTTGTAGCTATATTT |
| 182 | TCATTTGGGGTCCTCTTTTGAGGAACAAGTTTTCTTGTCGCGAGCTGA |
| 183 | AAAGGTGGCATCCTCTTTTGAGGAACAAGTTTTCTTGTTCAATTCTAC |
| 184 | TAATAGTAGTTCCTCTTTTGAGGAACAAGTTTTCTTGTAGCATTAACA |
| 185 | TCCAATAAATTCCTCTTTTGAGGAACAAGTTTTCTTGTCATACAGGCA |
| 186 | AGGCAAAGAATCCTCTTTTGAGGAACAAGTTTTCTTGTTTAGCAAAAT |
| 187 | GTACCAAAAACATTATGACCCTGTAATACTTTTGCGGG |
| 188 | AGAAGCCTTTATTTCAACGCAAGGATAAAAATTTTTAG |
| 189 | AACCCTCATATATTTTAAATGCAATGCCTGAGTAATGT |
| 190 | GTAGGTAAAGATTCAAAAGGGTGAGAAAGGCCGGAGAC |
| 191 | AGTCAAATCACCATCAATATGATATTCAACCGTTCTAG |
| 192 | CTGATAAATTAATGCCGGAGAGGGTAGCTATTTTTGAG |
| 193 | AGATCTACAAAGGCTATCAGGTCATTGCCTGAGAGTCT |
| 194 | GGAGCAAACAAGAGAATCGATGAACGGTAATCGTAAAA |
| 195 | CTAGCATGTCAATCATATGTACCCCGGTTGATAATCAG |
| 196 | AAAAGCCCCAAAAACAGGAAGATTGTATAAGCAAATAT |
| 197 | TTAAATTGTAAACGTTAATATTTTGTTAAAATTCGCAT |
| 198 | TAAATTTTTGTTAAATCAGCTCATTTTTTAACCAATAG |
| 199 | GAACGCCATCAAAAATAATTCGCGTCTGGCCTTCCTGT |
| 200 | AGCCAGCTTTCATCAACATTAAATGTGAGCGAGTAACA |
| 201 | ACCCGTCGGATTCTCCGTGGGAACAAACGGCGGATTGA |
| 202 | CCGTAATGGGATAGGTCACGTTGGTGTAGATGGGCGCA |
| 203 | TCGTAACCGTGCATCTGCCAGTTTGAGGGGACGACGAC |
| 204 | AGTATCGGCCTCAGGAAGATCGCACTCCAGCCAGCTTT |
| 205 | CCGGCACCGCTTCTGGTGCCGGAAACCAGGCAAAGCGC |
| 206 | CATTCGCCATTCAGGCTGCGCAACTGTTGGGAAGGGCG |

Continued on next page

Table F1 – continued from previous page

| Position | Sequence (5' → 3') |
|---|---|
| 207 | ATCGGTGCGGGCCTCTTCGCTATTACGCCAGCTGGCGA |
| 208 | AAGGGGGATGTGCTGCAAGGCGATTAAGTTGGGTAACG |
| 209 | CCAGGGTTTTCCCAGTCACGACGTTGTAAAACGACGGC |
| 210 | CAGTGCCAAGCTTGCATGCCTGCAGGTCGACTCTAGAGGATCTTTT |
